# Sequence Alignment Through the Looking Glass

**DOI:** 10.1101/256859

**Authors:** Raja Appuswamy, Jacques Fellay, Nimisha Chaturvedi

## Abstract

Rapid advances in sequencing technologies are producing genomic data on an unprecedented scale. The first, and often one of the most time consuming, step of genomic data analysis is sequence alignment, where sequenced reads must be aligned to a reference genome. Several years of research on alignment algorithms has led to the development of several state-of-the-art sequence aligners that can map tens of thousands of reads per second.

In this work, we answer the question “How do sequence aligners utilize modern processors?” We examine four state-of-the-art aligners running on an Intel processor and identify that all aligners leave the processor substantially underutilized. We perform an in-depth microarchitectural analysis to explore the interaction between aligner software and processor hardware. We identify bottlenecks that lead to processor underutilization and discuss the implications of our analysis on next-generation sequence aligner design.

## I. Introduction

The past few years have witnessed dramatic improvement in both cost and throughput of DNA sequencing technologies. Today, it is possible to sequence a single human genome at 30-fold coverage for as little as $1,000. With sequencing becoming more affordable, the amount of genomic data generated has been increasing at an alarming rate far outpacing Moore’s law [20]. This data deluge has the potential to pave way for the emerging field of personalized medicine and assist in detecting when genomic mutations predispose humans to certain diseases like cancer, autism, and aging.

However, to unlock the potential of genomic data, one needs scalable, efficient data analytics platforms and tools. The first and one of the most time-consuming steps in analyzing such data is sequence alignment–the task of determining the location in the reference genome that corresponds to each sequenced read. In the last decade, researchers have designed over 70 read mapping tools [8], each differing from another with respect to accuracy, sensitivity, specificity, and speed. Today, state-of-the-art read aligners can map tens of thousands of reads per second to the reference genome.

While all prior research focuses on improving performance, by reducing mapping time of individual reads, or scalability, by mapping more reads per second, there is no study that analyzes sequence aligners at the microarchitectural level. Such an analysis is important for several reasons.

First, it answers the following question–Do state-of-the-art aligners use modern processors efficiently? Microarchitectural analysis in other applications areas, like relational databases and data analytics platforms, showed that these applications do not utilize modern processors efficiently [1], [6]. Such analyses spurred research efforts to improve performance by redesigning data structures, or energy efficiency by using low-power processors that can match application requirements. In this work, we perform a similar analysis for sequence alignment.

Second, modern processors are complex pieces of hardware that use a variety of techniques to execute instructions faster. However, in order for such improvements to translate into tangible performance benefit, software must be optimized to avoid bottlenecks. Microarchitectural analysis reveals these bottlenecks by exposing harmful interaction between application software and processor hardware and helps in answering the following question–Will current software automatically benefit from microarchitectural improvements in the next-generation of hardware?

Third, the past few years have witnessed a rise in adoption of heterogeneous computing, as accelerators like General-Purpose Graphics Processing Units (GPGPU) and Intel Xeon Phi are being increasingly adopted in several data-intensive application domains. Given that sequence alignment is a complex, multi-stage process, several researchers have built aligners that execute some, or all stages of sequence alignment, on these accelerators [11], [18], [19], [21]. However, there has been no systematic analysis that explores the interaction between alignment stages and CPU microarchitecture to clearly identify which stages are more suited to the GPGPU than the CPU.

In this work, we present, to our knowledge, the first microarchitectural analysis of four state-of-the-art sequence aligners. We show that despite a decade of research and optimized implementations, modern aligners substantially underutilize processor resources as the processor is stalled in more than 50% of execution cycles without doing useful work. We provide an in-depth breakdown of stall cycles to identify hardware components in the processor pipeline and algorithmic components in sequence alignment software that contribute to these stalls. We discuss the implications of our analysis on the design of next-generation of sequence aligners and suggest directions for further research.

The rest of this paper is organized as follows. In Section II, we provide a brief overview of alignment techniques and processor microarchitecture. In Section III, we describe the hardware and software setup we use for this analysis. We present a global microarchitectural analysis of aligners in Section IV and a stage-by-stage analysis of one the aligners in Section V. Finally, we present the implications of our analysis in Section VI and conclude in Section VII.

## II. Background

In this section, we will provide a brief overview of sequence alignment techniques and modern processor microarchitecture to set the context for this work. We refer the reader to prior work for an in-depth algorithmic survey of sequence alignment algorithms and experimental analysis of aligners [5], [8]–[10].

### A. Sequence alignment

Modern Next-Generation Sequencing (NGS) technologies produce millions of short string sequences, referred to as reads, with each sequence corresponding a portion of the DNA. The first step in the analysis of this NGS data, referred to as sequence alignment, is to determine the location in the genome that corresponds to each of these short reads. Thus, sequence alignment is essentially a string matching problem where given a string G (reference genome), and a set of substrings R (reads), the origin of each read r ∈ R must be identified in G. However, due to sequencing errors or due to differences between the reference genome and the sequenced organism, a read might not exactly match its corresponding location in the reference genome. Thus, an aligner has to perform approximate string matching that is tolerant to mismatches, insertions, and deletions.

Since a read could potentially align at each one of the 3 billion locations in the reference, brute force search of each possible alignment is infeasible even for a single read. Thus, all modern aligners build an index over the reference and use the index to quickly narrow down the search space of potential locations. Aligners can be broadly classified into two types based on the indexing technique used, namely, seed-and-extend (SE) aligners or Burrows-Wheeler-Transform (BWT)-based aligners.

SE aligners typically use a hashtable to index the reference genome and store the contents and the occurrence locations of short string sequences, also referred to as seeds or k-mers, in a hash table. Each read is processed in three steps. First, seeds are extracted from the read and the hash table is used to look up potential mapping locations in the reference genome. Second, filtering techniques are used to reduce the number of potential locations where further extension must be performed. Third, the entire read is aligned at each of potential reference location using an approximate string matching algorithm like Needleman-Wunsch or Smith-Waterman.

BWT-based aligners align reads against a suffix array built using the reference genome. As suffix arrays are memory intensive, all of these aligners use a space-optimized data structure called FM-index [7] that uses a compressed string representation called Burrows-Wheeler Transform [4] to index the reference genome and enable fast exact string matching. As the FM-Index itself does not allow approximate string matching, BWT-based aligners use an algorithmic technique called backtracking that tries to insert errors at various positions in the read as it is matched while traversing the FM-Index. As the cost of backtracking increases exponentially, BWT-based aligners also use heuristics to prune the search space.

BWT-based alignment, introduced by Bowtie [12] and BWA [16], has been the most popular technique for aligning short reads. The low error rate of NGS technologies and a low divergence between reference and sequenced organisms allowed aligners like trie-based Bowtie and BWA to traverse the search space for very short reads 10 to 100 times faster than hash-based aligners. However, with read lengths increasing to 100-150 base pairs, backtracking emerged as the bottleneck especially if one needs to tolerate a larger number of errors. Thus, newer variants of these aligners, like Bowtie2 [2] and BWA-MEM [15] have also reverted back to using the seed-and-extend technique.

### B. Modern processor microarchitecture

In order to understand how state-of-the-art sequence aligners utilize modern processors, we need to examine how well are the microarchitectural resources of a processor used by the aligner software. Modern processors are typically multi-core in nature and contain several of processing cores. State-of-the-art aligners exploit the task parallelism offered by multi-core processors to scale sequence alignment by using multiple threads, one per core, where each thread aligns a disjoin set of input reads. While recent studies have focused on software issues that prevent scalability on multi-core processors [13], [14], in this paper, our focus is on the utilization of a single processing core. Thus, in the rest of this paper, we will use the term processor and core interchangeably.

The microarchitectural pipeline of a modern high-performance processor is quite complex. Figure 1 shows a simplified view of the microarchitectural components of a processing core. The pipeline of a processor is divided conceptually into two halves, the Front-end (FE) and the Back-end (BE). The FE is responsible for fetching the program code corresponding to the Instruction Set Architecture and decoding them into one or more low-level hardware operations called micro-operations (*µ*Ops). Once decoded, these *µ*Ops are queued for execution by the BE. Before a *µ*Op can be executed, all necessary data operands must be fetched from memory if necessary. The BE scheduler is responsible for monitoring when a *µ*Op’s data operands are available. Once ready, the scheduler associates the *µ*Op with a port depending on its intended execution purpose. For instance, a *µ*Op corresponding to an arithmetic operation would be associated with a port between 0 and 2, while a *µ*Op associated with loading data from memory would be associated a port between 3 and 5 in Figure 1. When an execution unit is available, the port dispatches the queued *µ*Op and executes it. Execution units, labelled as ALU, Divide, Mul, etcetera, in Figure 1, are the work horses that perform various operations like memory loads and stores, addition, multiplication, division, etcetera. The FE and BE are also equipped with both on-chip and off-chip cache memory, shown as L1/L2/L3 instruction/data caches in Figure 1, to avoid long-latency DRAM accesses by buffering data and instructions.

**Fig. 1:**
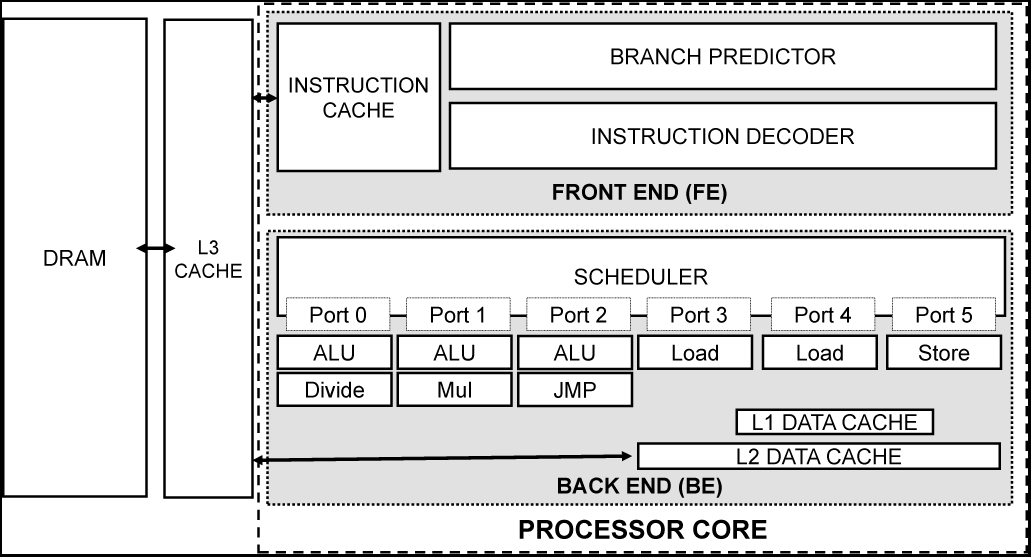
Simplified microarchitectural depiction of modern processors.

The completion of a *µ*Op’s execution is called *retirement*. When a *µ*Op is retired, its results are committed back to the architectural state by updating CPU registers or writing back to memory. The Front-end of the pipeline on recent Intel microarchitectures can allocate four *µ*Ops per clock cycle, while the Back-end can retire four *µ*Ops per clock cycle. Thus, in each clock cycle, modern Intel processors can potentially execute four instructions simultaneously. During instruction execution, most *µ*Ops pass completely through the pipeline and retire. But sometimes, a *µ*Op that is not be able to complete immediately might delay, or *stall*, the pipeline.

Intel’s Top-Down Analysis Methodology [22] classifies stalls into three major types, namely *Front-end* stalls, *Speculation* stalls, and *Back-end* stalls. During a cycle, if the BE is ready to execute a *µ*Op but the FE is unable to queue it for execution, the stall is classified as a FE stall. A typical reason for FE stalls is instruction cache misses caused by large instruction footprint corresponding to a complex code base. Disk-based relational database engines are known to suffer from such stalls [1].

Modern processors use speculative execution to improve instruction throughput. Conditional execution in programs, like if–else blocks, get translated into branch instructions that decide control flow depending on predicates in the conditional statement. Before the processor has to execute a branch instruction, the predicate value has to be determined. Instead of waiting until a branch instruction’s predicate is resolved, the Branch Predictor component in the FE implements an algorithm that guesses the predicate and fetches the appropriate instruction stream. If the guess is correct, the execution continues normally, and if it is wrong, the pipeline is flushed, and the correct instruction stream is fetched and executed. Such flushing creates pipeline stalls and these stalls caused by incorrect branch prediction are classified as Speculation stalls.

In the case where the FE has a *µ*Op ready but the BE is not ready to handle it, the processor is stalled on the BE. BE stalls can be further classified into *Memory stalls* and *Core stalls*. As mentioned earlier, a *µ*Op can be executed only if its data operands are available. Memory stalls are caused by the processor having to wait for such data operands to be fetched from the cache or from memory. Core stalls, in contrast, are caused by a less-than-optimal use of the available execution units during each cycle. This can happen due to contention for resources. For instance, if the code contains instructions that result in several *µ*Ops being associated with a few ports, then queue for those ports become full. Thus, the scheduler can no longer associate any further *µ*Ops with those ports until the queue shrinks. Similarly, if the code contains several divide instructions in a row, they will compete for the few divide execution units resulting in resource conflicts.

## III. Experimental Setup

In this section, we describe the hardware and software setup we use in this analysis and outline the experimental methodology.

### A. Hardware–software setup

All experiments are conducted on a server running RHEL 7.2, equipped with a 12-core Intel Xeon E5-2650L v3 CPU and 256GB RAM. We analyze four state-of-the-art sequence aligners, namely, *BWA-MEM* [15] and *Bowtie2* [2], *Snap* [23], and *FSVA* [17]. We chose *Bowtie2* and *BWA-MEM* as they are the most popular short-read aligners that use a BWT-based reference index. We chose *Snap* as a representative of a new breed of hashtable-based aligners that exploit the increasing read lengths to improve performance without sacrificing accuracy. We chose *FSVA*, as it is a state-of-the-art aligner built for cohort studies that explicitly trades off accuracy for fast single-threaded performance.

As described in Section II, all these aligners use the seed-and-extend technique to perform fast alignment of reads. However, these aligners differ dramatically with respect to the actual methodology used for seed selection, filtration, and extension. As a result, these aligners occupy different points in the performance–accuracy dimensions. Our goal in this analysis is to not to perform a side-by-side analysis of execution time or accuracy of various aligners. The optimal aligner choice is a complex analysis topic covered by prior research [5], [8]–[10] as there is a delicate balance between accuracy and performance that must be met depending on the expected usage. As our focus in this paper is on microarchitectural analysis of sequence aligners, we run each aligner only in the single threaded mode. Thus, we report the execution time and accuracy results only for completeness.

### B. Experimental methodology

We use both synthetic and real datasets for evaluating each system. The synthetic dataset is generated by using wgsim. We generate two datasets with one million reads of length 150 bp each, one with the default rate of indels and one without any indels or substitutions. *BWA-MEM*, *Bowtie2*, and *Snap* can be configured via command-line parameters to trade off alignment accuracy for improved performance. Thus, we use the two simulated datasets to perform a sensitivity analysis to the presence of indels. More specifically, for the indel-free case, we configure aligners for maximum performance by (i) increasing the reseeding parameter (-r) of *BWA-MEM* from 1.5 (default) to 10 (no further improvement beyond this on our hardware and dataset), (ii) using the “–very-fast” configuration option of *Bowtie2*, and (iii) setting the MaxDist parameter (-d) to zero for *Snap*. For the with-indel dataset, we run all aligners using default configuration parameters, thus trading off performance for accuracy. It is important to note that our goal is not to systematically explore the entire parameter space, but rather identify trends in CPU utilization at extreme points in the performance–accuracy spectrum. For the real dataset, we use a paired-end read (sample NA12878/09252015) obtained from the public Genome-In-A-Bottle (GIAB) dataset [24]. Similar to the simulated with-indel case, we run all aligners using default configuration parameters for the GIAB dataset.

We use Intel VTune for profiling each system. Before profiling, each aligner is run once to warm up the file system cache and ensure that the necessary indices and input data are memory resident. Then, we profile each aligner by executing it for 30 seconds, to warm up the instruction cache, and then attaching VTune to the target thread for another 30 seconds. We repeat the analysis three times and report only the median values as the variation across runs was less than 10%.

## IV. Analysis Results

In this section, we present our analysis of the four sequence aligners with respect to processor utilization.

### A. Indel-free dataset analysis

Modern processors use an array of techniques like pipelining, out-of-order execution, speculation, and instruction prefetching to improve single-threaded performance. As a result, modern processors can retire multiple instructions per clock cycle as described in Section II. The efficiency of a software is determined by the metric Instructions Per Cycle (IPC), which determines the number of machine instructions executed and retired by the processor in each clock cycle. The Intel processor we use in this analysis can retire four instructions per cycle. Thus, in the ideal case, the IPC value of a sequence aligner should be four.

#### IPC analysis

Table I shows the single-threaded execution time of aligners and Fig 2 shows the IPC values for the four aligners under the indel-free simulated dataset. Clearly, the processor remains substantially underutilized across aligners as the worst-case IPC is lower than 25% (for *BWA-MEM*), and the best-case IPC is around 50% (for *Bowtie2*), of the theoretically achievable maximum.

**TABLE I:** Execution time in seconds of aligners under different datasets

| Aligner | Simulated Indel-Free | Simulated With-Indel | GIAB |
| --- | --- | --- | --- |
| bowtie2 | 176 | 437 | 3126 |
| bwa | 127 | 458 | 2919 |
| snap | 57 | 112 | 717 |
| fsva | 55 | 80 | 288 |

**Fig. 2:**
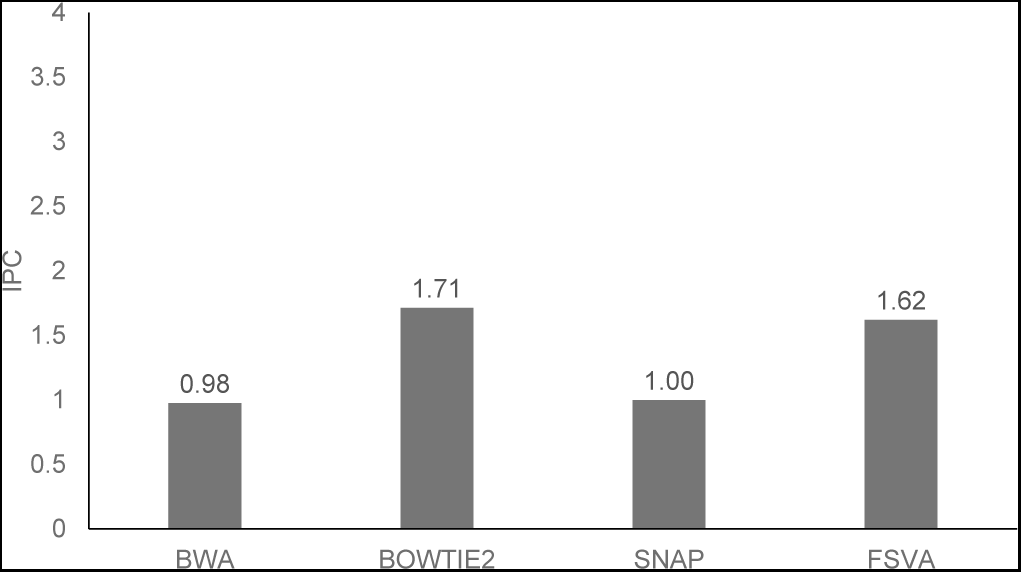
IPC for indel-free dataset

#### Execution cycle breakdown

In order to further understand why observed IPC values are lower than the theoretical maximum and explain differences in IPC across aligners, we need to analyze the processor activity on a finer microarchitectural level. Figure 3 shows the breakdown of execution cycles for each aligner into retiring and stalled for the indel-free dataset. We see that the processor is stalled in nearly 60-80% of cycles. These stalls result in the low IPC values observed earlier.

**Fig. 3:**
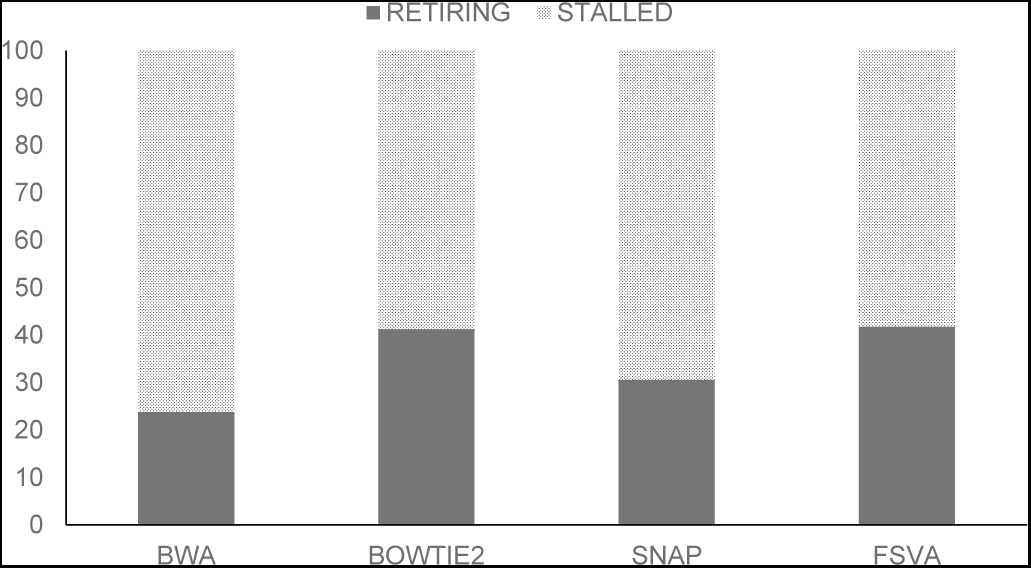
Execution cycle breakdown for indel-free dataset

#### Stall-cycle breakdown

Given that a substantial fraction of cycles are stall cycles, the next step is to identify the root cause of these stall cycles. Figure 4 shows the stall cycle breakdown for each aligner under the indel-free dataset. Clearly, the dominating source of stalls is the Back-end which accounts for 60% to 80% of all stall cycles. Figure 5 breaks down the Back-end stalls further into memory or core bound. Memory stalls account for a 60% to 80% of Back-end stalls across all aligners. This indicates that the processor is stalled waiting for data.

**Fig. 4:**
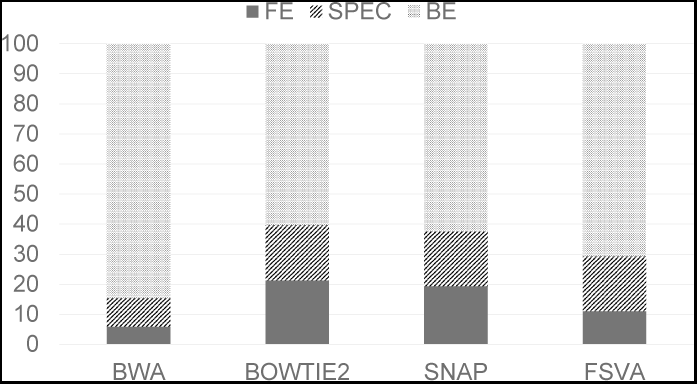
Stall cycle breakdown for indel-free dataset

**Fig. 5:**
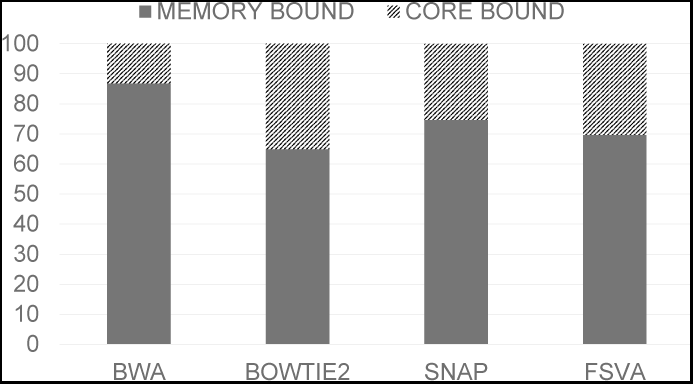
Back-end stall breakdown for indel-free dataset

#### Memory stalls analysis

Given that memory stalls dominate the Back-end across all aligners, it is important to know if these stalls are due to cache-resident data or DRAM-resident data. The Intel processor we use in this study has a three-level caching hierarchy, with a 32KB L1 cache, 256KB L2 cache, and 30MB Last-level cache. In general, software optimizations attempt to move data closer to the processor so that critical data structures are L1-cache resident. Thus, a memory stall at a lower cache level typically indicates an optimization opportunity where data structure redesign can improve utilization. However, if memory stalls are due to DRAM-resident data, this is typically due to cache-unfriendly random data access pattern which is harder to optimize.

Figure 6 breaks down memory stalls into load and store stalls. Load stalls are further decomposed based on the location that contributes to the stall (L1, L2, L3 cache, or DRAM). Clearly, between 70–90% of memory stalls are DRAM stalls indicating that the processor is waiting for long-latency memory accesses from DRAM.

**Fig. 6:**
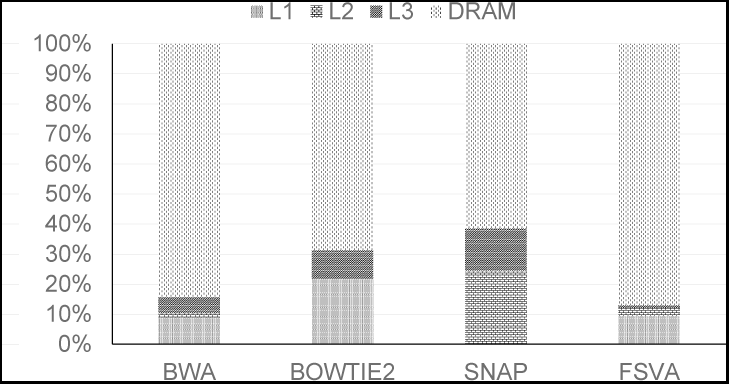
Memory stall breakdown for indel-free dataset

#### Insights

All aligners substantially underutilize the processor, as over 50% of execution cycles are spent on stalls. The processor is stalled as the Back-end is blocked on long-latency DRAM accesses waiting for data.

### B. Analysis of dataset with indels

Having analyzed the microarchitectural behavior of aligners under the indel-free dataset, we now present our results using the with-indel dataset. Our goal is to understand if the presence of indels, and the associated changes in aligner parameters to improve accuracy, results in a different microarchitectural behavior compared to the indel-free case where aligners were configured for peak performance.

#### IPC analysis

Table I shows the single-threaded execution time of aligners under the with-indel simulated dataset. Comparing the execution times between indel-free and with-indel datasets in Table I, we see that the execution time of all aligners increases in the presence of indels. This is expected given that the indels trigger expensive approximate alignment algorithms. Figure 12 shows the ROC curve generated by wgsim eval. The figure plots the number of correct alignments against the number of incorrect alignments as mapping quality decreases from left to right. Clearly, *BWA-MEM*, and *Bowtie2* provide the most accurate alignment for this dataset followed by *Snap* and *FSVA*. While our goal is not to provide a side-by-side comparison of aligners, these results clearly indicate that these aligners occupy different points in the performance–accuracy spectrum.

Figure 7 shows the IPC values under the with-indel dataset. Comparing Figures 2, 7, we see that the IPC value of aligners under the with-indel dataset is higher than the indel-free dataset.

**Fig. 7:**
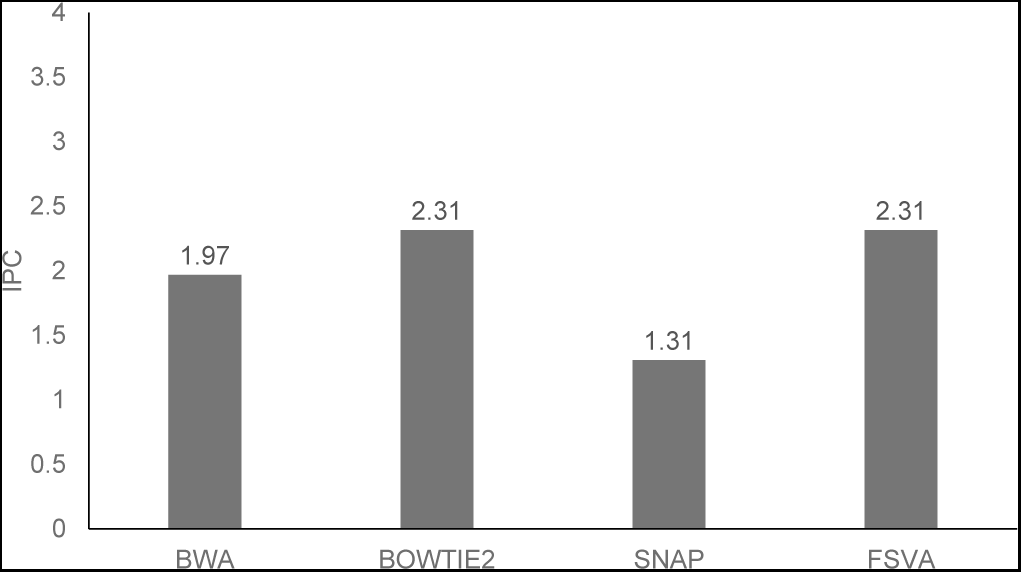
IPC for with-indel dataset

#### Execution cycle breakdown

Figure 8 shows the break-down of execution cycles for each aligner into retiring and stalled for the with-indel datasets. Similar to the indel-free dataset (Figure 3), we see that even in the best case, the processor is still stalled in nearly 50% of cycles. However, the processor retires more instructions when the dataset has indels compared to a dataset without indels. This directly translates into corresponding difference in IPC observed earlier between the two datasets.

**Fig. 8:**
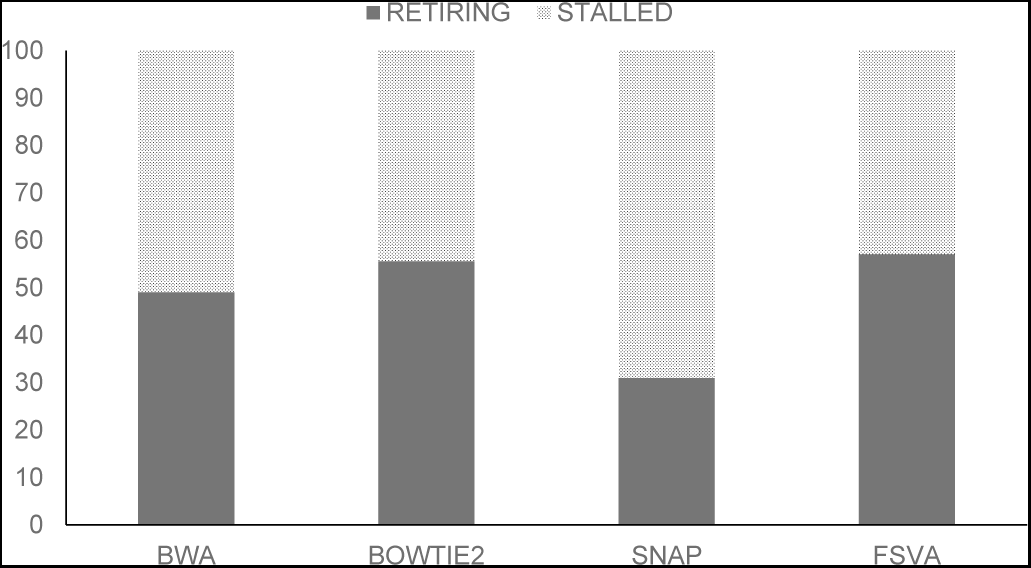
Execution cycle breakdown for with-indel dataset

#### Stall cycle breakdown

Figure 9 shows the stall cycle breakdown for each aligner under the with-indel dataset. We can make two important observations. First, under *Bowtie2*, *BWA-MEM*, and *FSVA*, the dominating source of stalls is still the Back-end which accounts for 60% to 70% of all stall cycles. However, comparing this with the indel-free case (Figure 4), we see that the contribution of speculation stalls increases in the presence of indels. Second, unlike the marginal increase in speculation stalls under *BWA-MEM*, *Bowtie2*, and *FSVA*, we see that speculation emerges as the dominating source of stalls under *Snap* accounting for nearly 45% of stall cycles. The Back-end contributes to 35% of stalls with *Snap*.

**Fig. 9:**
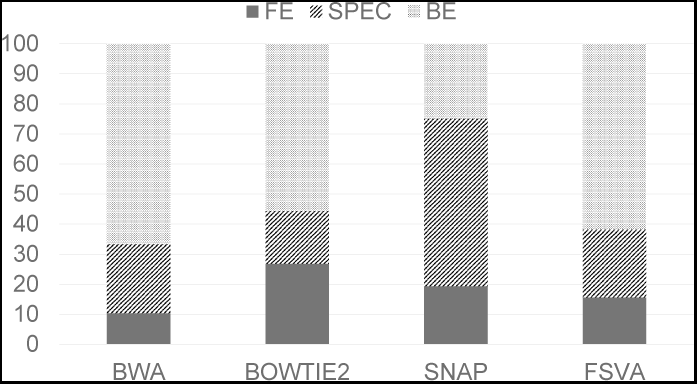
Stall cycle breakdown for with-indel dataset

Figure 10 breaks down the Back-end stalls further into memory or core bound for *Bowtie2*, *BWA-MEM*, and *FSVA*. We see that while memory stalls still account for over 40% of Back-end stalls, they are no longer the only dominating source, as core stalls account for as much as 60% of stalls under some aligners. This contrasts sharply with the indel-free case (Figure 5), where memory stalls overshadow core stalls. These results show that aligners choose different code paths for dealing with indel-free and with-indel cases as expected. Further, the code path executed under the with-indel dataset stresses the microarchitecture in a different way compared to the code path executed under the indel-free case.

**Fig. 10:**
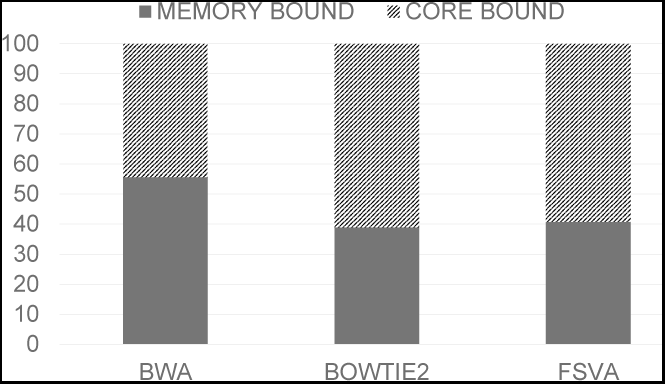
Back-end stall breakdown for with-indel dataset

#### Memory stalls analysis

Given that memory stalls still account for atleast 40% across aligners, Figure 11 breaks down memory stalls across various cache levels and DRAM under the with-indel dataset. Comparing Figures 6, 11, we see that under *BWA-MEM* and *FSVA*, long-latency DRAM stalls continue to dominate and contribute to over 80% of all memory stalls in both indel-free and with-indel datasets.

**Fig. 11:**
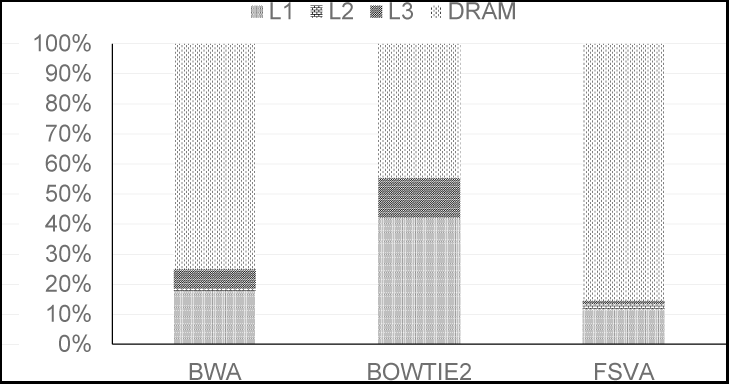
Memory stall breakdown for with-indel dataset

**Fig. 12:**
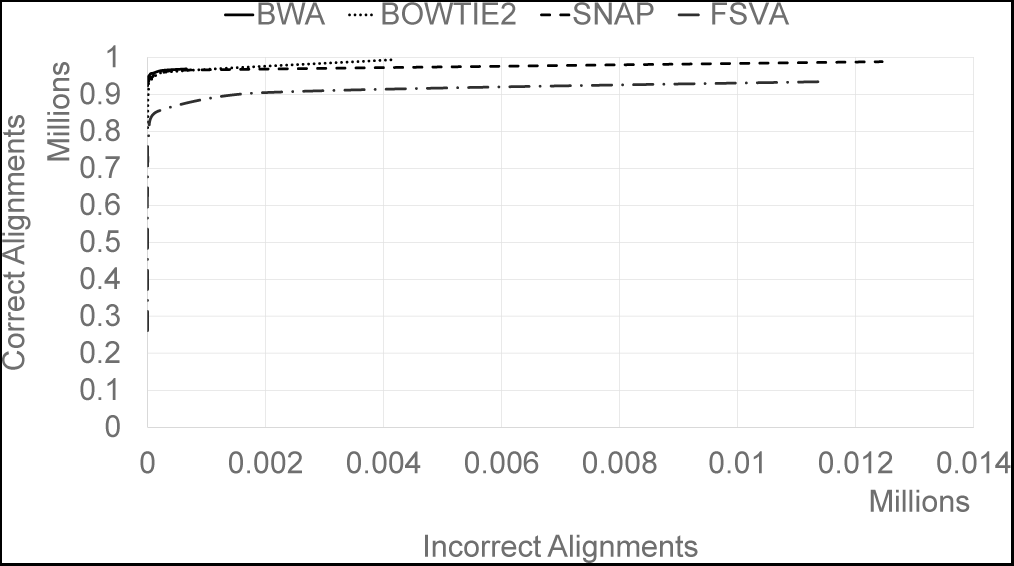
ROC curve for with-indel dataset

#### Insights

While processor utilization improves in the presence of indels across all aligners, the processor still spends majority of its cycles in the stalled state. However, memory stalls are no longer the majority contributor, as aligners also suffer from speculation and core stalls.

### C. GIAB dataset analysis

So far, we have presented our analysis based on the simulated datasets. The microarchitectural behavior of aligners under the GIAB dataset is very similar to the simulated, with-indel dataset.

Table I shows the single-threaded execution time of aligners under the GIAB dataset. Figures 13, 14 show the IPC and execution cycle breakdown. Comparing this with Figure 8, we see similar trends between the with-indel simulated dataset and the GIAB dataset as the processor is stalled in 50-70% of execution cycles.

**Fig. 13:**
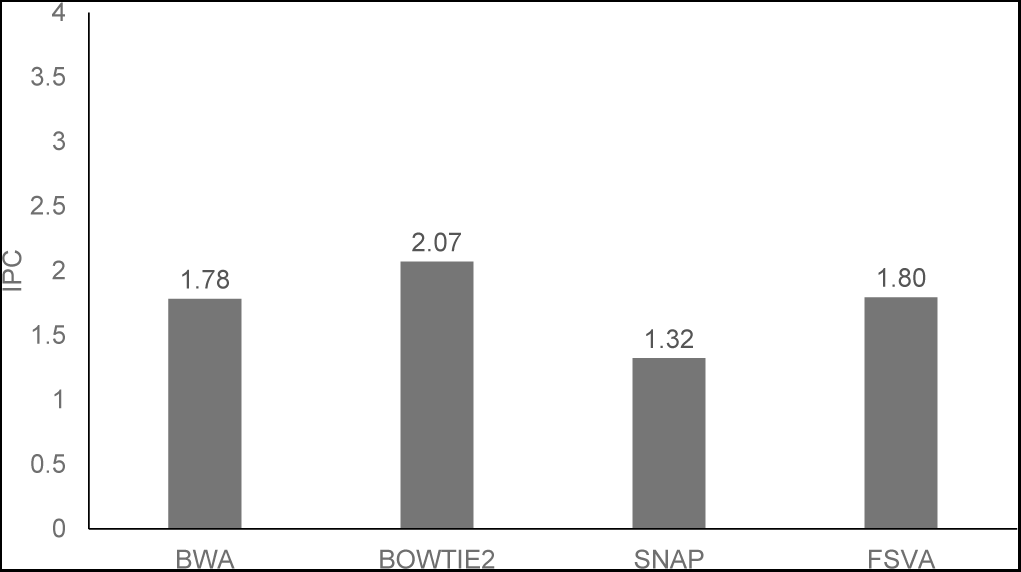
IPC under GIAB dataset

**Fig. 14:**
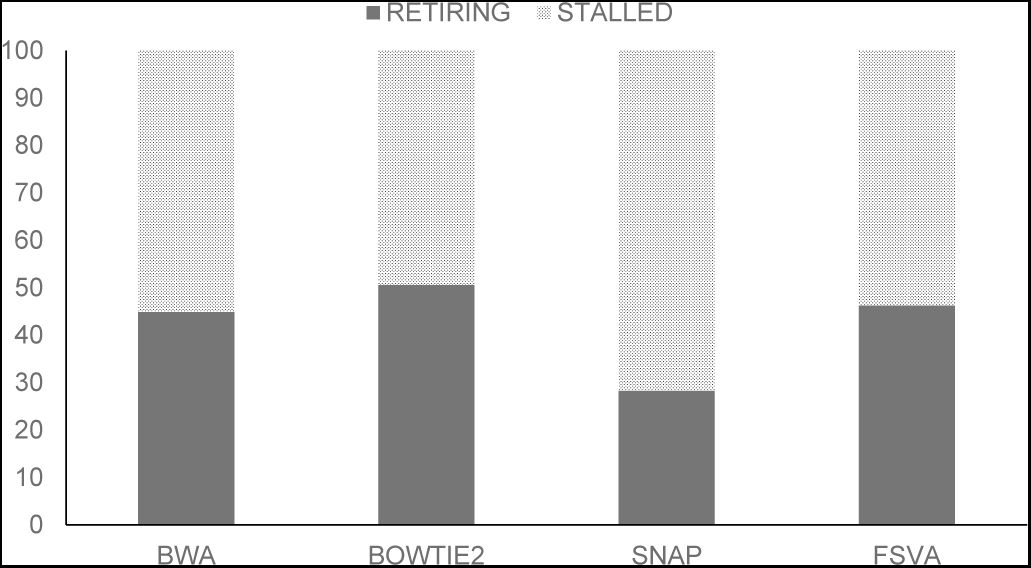
Execution cycle breakdown under GIAB dataset

Figure 15 breaks down the stall cycles into various components. Similar to the simulated dataset (Figure 8), Back-end stalls account for 60–70% of stall cycles under *Bowtie2*, *BWA-MEM*, and *FSVA*. Speculation stalls are the dominating source of stall cycles under *Snap*.

**Fig. 15:**
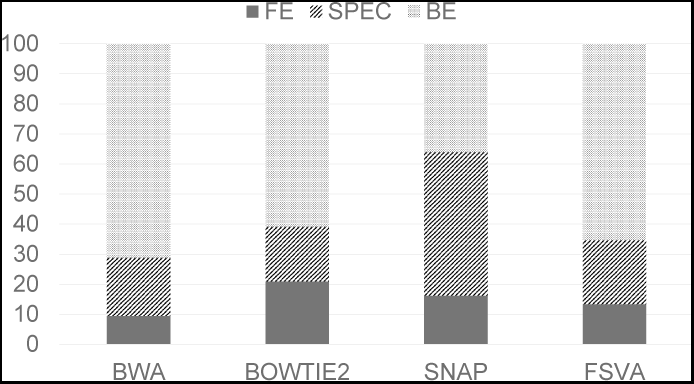
Stall cycle breakdown for GIAB dataset

Among aligners bottlenecked on Back-end stalls, Figure 16 shows that memory stalls are the dominating factor. Figure 17 shows that long-latency DRAM stalls are the main contributors for memory stalls.

**Fig. 16:**
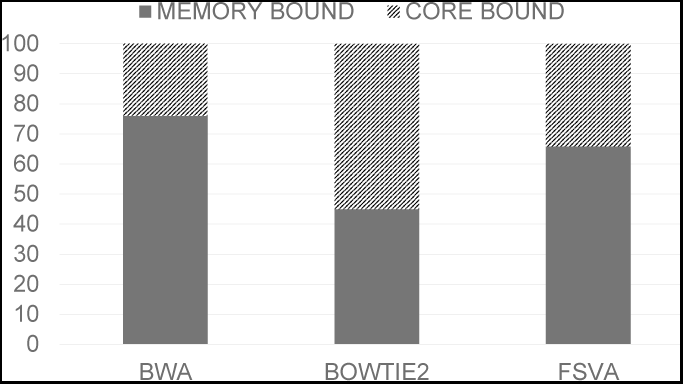
Back-end stall breakdown for GIAB dataset

**Fig. 17:**
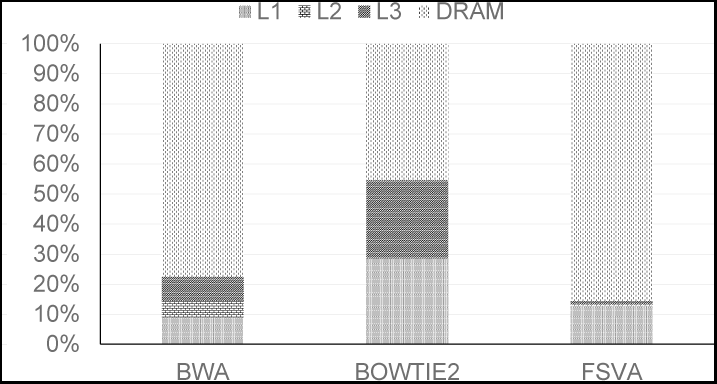
Memory stall breakdown for GIAB dataset

## V. Per-stage analysis

The analysis presented so far answers some questions–how do aligners utilize the processor? what causes processors to be stalled? But, it also raises other questions–why does the presence of indels change the microarchitectural behavior? To answer this question, we need to perform a stage-by-stage analysis of sequence alignment.

All aligners we have considered in this study work by considering one read at a time. Each aligner processes each read using three distinct steps. In the first stage, seeds are extracted from each read and used to lookup the index to retrieve candidate locations. The second stage is the filtration stage where heuristics and theoretical lower bounds are used to reduce the number of candidate locations that must be examined. The third stage is the extension stage where the entire read is aligned with each candidate location to identify the best match. Once a read has been processed by all the three stages, the aligner writes the alignment information to the output file and moves to the next read.

The analysis results we have presented so far are from an inter-stage analysis that spans across all stages. In this section, we present intra-stage analysis where we explore processor utilization within each stage to identify which stages account for stall cycles so that we can answer the aforementioned question. In order to perform the intra-stage analysis, we modified *FSVA* so that data is processed in a stage-at-a-time fashion. We chose *FSVA* as it was the fastest aligner in our study and has a simpler code base due to its focus on single-threaded performance.

In our modified *FSVA*, which we henceforth refer to as *staged-FSVA*, all reads are first broken down into seeds and a hashtable lookup is performed to identify candidate locations. All such locations are saved in intermediate data structures together with metadata to keep track of the mapping between locations and reads. Once all reads are processed by the first stage, the output from the first stage is passed to the second stage. In this stage, the candidate locations are sorted and filtered using the seed-and-vote filtration approach used by FSVA [17]. The output of this stage are two candidate locations that are the two most voted locations for each read. Once these location pairs are identified for all reads, staged-FSVA proceeds to the third stage where Smith-Waterman alignment is used to perform approximate alignment.

By using a stage-at-a-time execution approach, staged-FSVA makes it possible for us to accurately measure and profile each stage independently.

### A. IPC and execution cycle breakdown

Table II shows a breakdown of execution time of each stage of staged-FSVA under the with-indel simulated dataset. As expected, the Smith-Waterman extension stage dominates overall execution time. Figure 18 shows the IPC values for each phase of staged-FSVA. While the filtration and extension stages have IPC values of 2.5 and 2.9, the seeding and hashtable lookup stage has only an IPC value of 0.5. This highlights that the main bottleneck leading to low IPC count, and hence poor processor utilization, is the first stage.

**TABLE II:** Execution time in seconds of the three main alignment stages and rest (loading reference, I/O, writing out SAM file, etcetera) of FSVA

| Stage | Time (secs) |
| --- | --- |
| Seeding and lookup | 14 |
| Filtration and voting | 11 |
| Extension | 35 |
| Misc. | 22 |

**Fig. 18:**
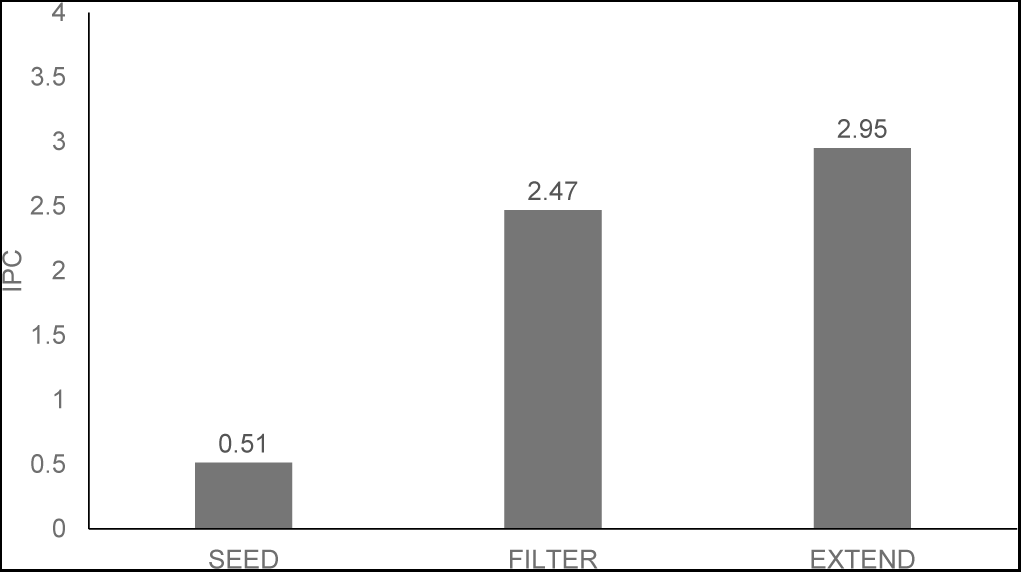
Per-phase IPC under simulated with-indel dataset

Figure 19 shows the breakdown of execution cycles per stage for staged-FSVA. These results mirror the IPC values in Figure 18. In the first stage, the processor spends 87% of execution cycles in the stalled state, while retiring instructions only in 13% of cycles. In the second and third stages, this trend is reversed, as the processor spends 63% to 72% of cycles retiring instructions.

**Fig. 19:**
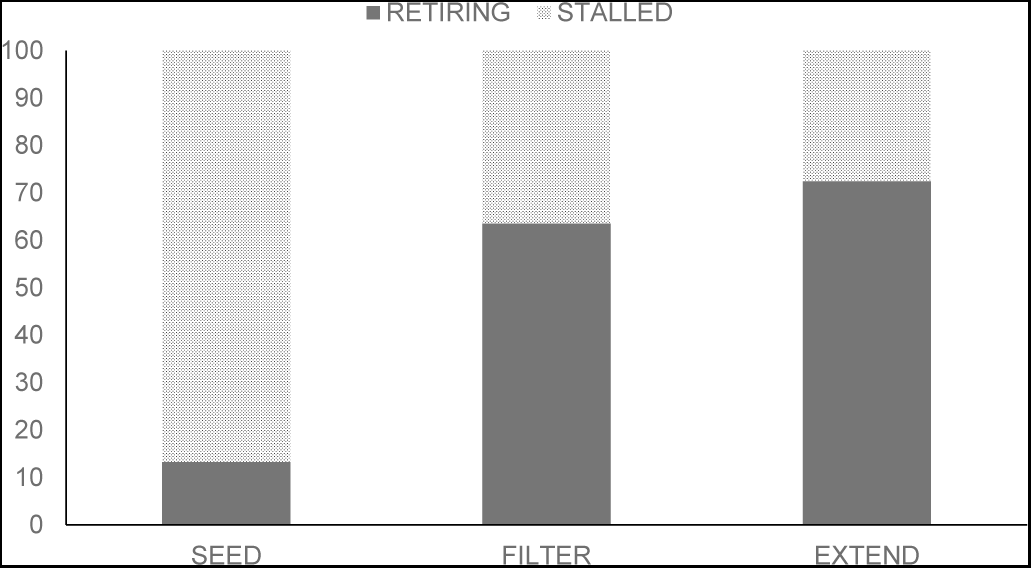
Per-phase execution cycle breakdown

### B. Stall cycle breakdown

Figure 20 shows the breakdown of stall cycles for each stage. Clearly, there is a dramatic difference between stages. In the seeding and lookup stage, Back-end stalls account for 90% of stall cycles. Figure 21 shows the breakdown of Back-end stalls and as can be seen, memory stalls contribute to over 90% of the Back-end stalls in the seeding stage. Figure 22 shows the memory stall breakdown. Clearly, long-latency DRAM access dominate this category and account for 93% of all memory stalls in the seeding stage.

**Fig. 20:**
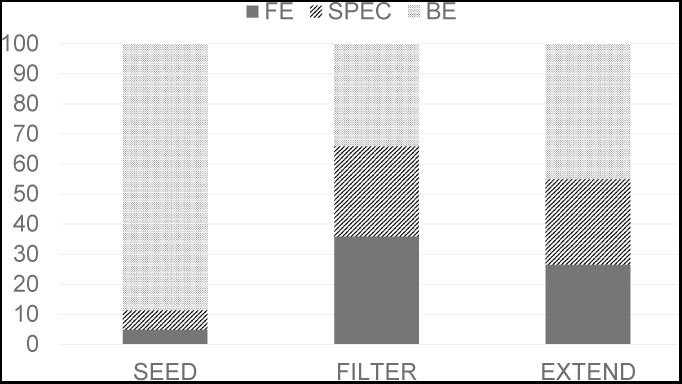
Per-phase stall cycle breakdown for with-indel dataset

**Fig. 21:**
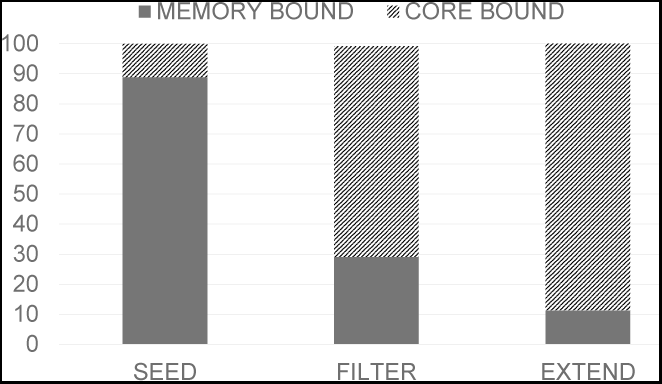
Per-phase Back-end stall breakdown for with-indel dataset

**Fig. 22:**
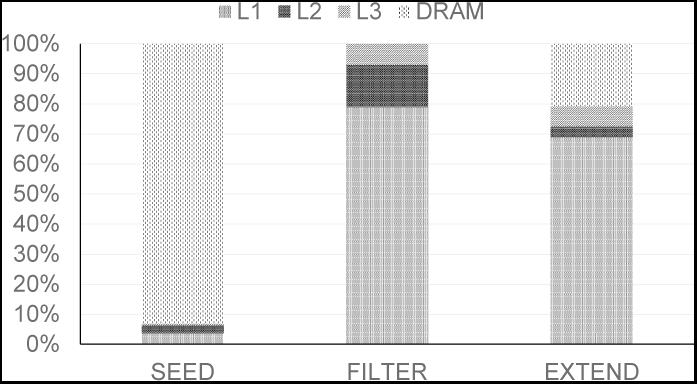
Per-phase memory stall breakdown for with-indel dataset

During this first stage of alignment, seeds of length 31 bp are extracted from each read, hashed, and used to probe the hashtable to determine coordinates in the reference. As each seed potentially hashes to a completely random value in a 4GB range, there is little spatial locality in this workload. Thus, hashtable lookups do not benefit from processor caches, the corresponding memory load instructions result in long-latency DRAM accesses. As instructions that follow these memory loads are dependent on the load, the pipeline is stalled and the processor idles waiting for data to arrive from memory.

Unlike the seeding stage, the filtration stage is not entirely bottlenecked on the backend. During filtration, Front-end, speculation, and Back-end stalls contribute equally to stall cycles as shown in Figure 21. *FSVA* uses a seed-and-vote-based filtration scheme where all coordinates gathered are used to vote for candidate locations where the read must be aligned. This voting is accomplished by sorting the coordinates determined in the seeding stage, eliminating duplicates and low-frequency locations, and identifying the top two locations with most votes. The branching logic in the sorting algorithm results in speculation stalls. Mispredicted branches add delay to the Front-End as it has to fetch operations from corrected path, resulting in Front-end stalls. The Back-end stall break-down shown in Figure 21 shows that core stalls are the major (71%) contributor to Back-end stalls. Figure 22 shows that the remaining 29% contribution from memory stalls is due to L1 cache accesses and not due to DRAM access. Thus, computation rather than memory access is the bottleneck in the filtration stage.

The extension stages is similar to the filtration stage microarchitecturally. As we already mentioned, the processor is stalled in only 30% of cycles (Figure 19). As shown in Figure 20, Back-end stalls account for 40% of execution cycles while Front-end and speculation account evenly for the other 60%. While Back-end stalls are relatively higher in the extension stage compared to the filtration stage, these stalls are once again due to core stalls, which contribute to nearly 90% of Back-end stalls, rather than memory stalls as shown in Figure 21. During the extension stage, *FSVA* uses the Smith Waterman dynamic programming algorithm for aligning reads to the reference. As the amount of data that the algorithm operates on fits easily in the processor cache, there are very few data misses. However, the complex control flow and branching logic in the dynamic programming algorithm creates speculation and core stalls.

#### Insights summary

There is a clear dichotomy between various stages of sequence alignment with respect to microarchitectural utilization. The seeding stage exhibits very low processor utilization with memory stalls in the Back-end contributing to majority of processor stall cycles. The filtration and extension stage, in contrast, have much better utilization, and are bottlenecked on Front-end speculation stalls and Back-end core stalls.

## VI. IMPLICATIONS

In this section, we will discuss the implications of our findings on the design of next-generation of sequence aligners. We will consider three dimensions, namely, performance, scalability, and energy efficiency.

### A. Performance implications

Even though modern sequence aligners are able to map reads to reference at a very high throughput, our analysis reveals that they still substantially underutilize the processor. Irrespective of the aligner used, the processor remains stalled over 50% of time suggesting that there is room for further improvement. Our analysis also showed that different stages of sequence alignment behave differently with respect to processor utilization. The seeding stage has the worst utilization of all stages as memory stalls caused by hashtable lookups dominate execution cycles. The filtration and extension stages, in contrast, utilize the processor better and do not experience such memory stalls.

Given this dichotomy between stages, it is clear that the two stages should be optimized differently. Given that the filtration and extension stages are bottlenecked on resource conflicts (core stalls), using “beefier” processors with more execution units, or using software techniques like vectorization to feed more data to existing units, should both assist in improving performance of these two stages. However, given that the seeding stage is bottlenecked on memory stalls, simply using faster processors will only result in an increase in stall cycles instead of improved performance. The solution is to use latency-hiding techniques for masking the overhead of memory accesses. Thus, techniques like software prefetching, simultaneous multithreading, can be explored further to ensure that the processor continues to retire instructions corresponding to one read while waiting for data to arrive from memory for another read.

### B. Scalability implications

Modern sequencing technologies produce millions of reads in a single run. Given that each read is independent of other reads, scaling sequence alignment is an embarrassingly parallel problem as each read can be assigned to a different processing thread. Given that modern servers are equipped with multicore processors, state-of-the-art aligners have started exploiting the thread-level parallelism of these processors to scale alignment. However, given that a single processor is stalled 50% of cycles, and given that various stages of alignment differ in their usage of processor resources, such an approach of using homogeneous multiprocessing where processors are identical to one another will only aggravate underutilization.

A promising alternative is to consider the use of heterogeneous parallelism using accelerators like Xeon Phi or GPGPUs. However, although GPGPUs provide massive thread-level parallelism with thousands of CUDA cores, research has shown that current GPGPU-based aligners provide only around 2–3 improvement compared to CPU-based aligners [19], [21]. Unlike CPUs which excel at task parallelism, GPGPUs excel at data parallelism. Sequence alignment, however, lends itself naturally to task parallelism rather than data parallelism due to the complex branching logic used in dynamic programming-based extension algorithms. Thus, current GPGPU-based aligners that attempt to execute the extension stage on GPGPUs suffer from scalability limitations due to warp divergence caused by branching logic.

Our analysis suggests a natural division between CPUs and GPGPUs. CPUs are underutilized substantially during the seeding phase due to long-latency data misses. GPGPUs, in contrast, are capable of hiding long-latency accesses using hardware-assisted multithreading. Thus, GPGPUs are a natural fit for the seeding phase of sequence alignment. Similarly, filtration phase of sequence alignment uses sorting, duplicate removal, and candidate selection. As these operations are data parallel, they also likely to benefit by execution on GPGPUs. However, given that the extension stage uses complex branching logic, and given that CPU utilization is already high during this stage, it might be better to schedule it on CPUs instead of GPGPUs. Thus, further research is necessary to understand the pros and cons of such a design as opposed to a CPU-only or GPU-only aligner.

### C. Energy efficiency implications

With the advent of cloud computing, it has become increasingly more important for data analytics platforms to be energy proportional [3], meaning that they consume power proportional to the amount of work performed. Any program that results in the processor stalling for a substantial portion of time adversely impacts energy proportionality as the power consumed by the processor is not used for performing useful work. Given that the processor is stalled for over 50% of the execution cycles under sequence aligners, we believe that there is much work to be done in improving the energy efficiency of these tools.

One promising research direction would be to use the dichotomy between stages to implement sequence alignment on heterogeneous big-LITTLE processor architectures like ARM. As processors are equipped with both “beefy” cores and “wimpy” cores, a sequence aligner that uses the latter for the first stage and former for the latter two stages would consume much less power than one built for contemporary server-grade processors.

## VII. Conclusion

Sequence alignment is the first stage of genomic data analysis and a very well-studied problem. Decades of research on scalable, high-performance approximate string matching algorithms have led to the development of fast sequence aligners that can map thousands of reads per second. However, there is no work that analyzes the efficiency of sequence aligners with respect to processor utilization.

In this study, we presented the first microarchitectural analysis of four state-of-the-art aligners. Our analysis on simulated datasets as well GIAB data revealed that all aligners result in substantial underutilization as the processor remains stalled for 50%-70% of execution cycles. We identified Back-end memory stalls and speculation stalls as leading sources of inefficiency. To understand the source of these stalls, we also presented a stage-by-stage analysis which mapped memory stalls to the seeding stage and speculation stalls to the filtration and extension stages. This microarchitectural study shows an in-depth view of processor usage for one of the many steps in genomic data analysis pipeline and opens up new opportunities for extending this analysis to other phases as well. Given the growing popularity of heterogeneous parallelism, such microarchitectural studies will play an important role in determining the ideal processor type for each phase of the genomic data analysis pipeline.

